# Finding high-quality metal ion-centric regions across the worldwide Protein Data Bank

**DOI:** 10.1101/619809

**Authors:** Sen Yao, Hunter N.B. Moseley

## Abstract

As the number of macromolecular structures in the worldwide Protein Data Bank (wwPDB) continues to grow rapidly, more attention is being paid to the quality of its data, especially for use in aggregated structural and dynamics analyses. In this study, we systematically analyzed 3.5 Å regions around all metal ions across all PDB entries with supporting electron density maps available from the PDB in Europe. All resulting metal ion-centric regions were evaluated with respect to four quality-control criteria involving electron density resolution, atom occupancy, symmetry atom exclusion, and regional electron density discrepancy. The resulting list of metal binding sites passing all four criteria possess high regional structural quality and should be beneficial to a wide variety of downstream analyses. This study demonstrates an approach for the pan-PDB evaluation of metal binding site structural quality with respect to underlying x-ray crystallographic experimental data represented in available electron density maps of proteins. For non-crystallographers in particular, we hope to change the focus and discussion of structural quality from a global evaluation to a regional evaluation, since all structural entries in the wwPDB appear to have both regions of high and low structural quality.

## 1. Introduction

Metal ions are important components in biological processes, especially at the biochemical and cellular levels. An estimated 30 to 40 percent of proteins across the combined proteome of the biosphere binds at least one metal ion [1,2]. Protein metal binding is part of many biochemical mechanisms including signal transduction, enzyme catalysis, and protein structural integrity [3–5]. The local protein structure environment around bound metal ions can provide clues to the biochemical and cellular function of the binding [6–8] and how sequence-based structural changes modulates metal binding [9,10]. However, the quality of 3D protein structural data around metal binding sites can vary dramatically from structure to structure, and especially from region to region [8,11]. Therefore, when analyzing metal binding site structure and dynamics, the quality of utilized worldwide Protein Data Bank (wwPDB) [12] entries should be evaluated, especially in the metal binding site region [2,8,13]. The potential impact of crystallization artifacts on ligand binding affinities has already been demonstrated [14]. Also, current methods and tools for this regional evaluation around metal ions, focused only on the PDB structural entry itself, have proven useful for weeding out some metal binding sites with poor regional structural quality [13]. However, comparison to the raw electron density itself represents a gold standard of evaluation against experimental data that can demonstrate the reliability and usability of a given metal binding site region [8,15]. These comparisons of metal binding site structure to the underlying electron density data has been facilitated by structure factor deposition requirements of the wwPDB since 2008 and electron density maps made available previously by the Uppsala Electron Density Server [16] and now by the PDB in Europe (PDBe) [17]. But this electron density evaluation of regional structural quality has been a tedious process done by manual visual inspection, without objective metrics of quality. To alleviate these shortcomings in electron density evaluation, we have developed new analysis and evaluation methods in a Python package called pdb-eda [18] that facilitate the systematic quality control of protein structural regions of interest across large numbers of wwPDB entries and their corresponding electron density maps. In this study, we apply pdb-eda to a systematic electron density analysis of all metal binding sites containing a bound metal ion. This analysis provides an evaluation of the quantity of metal binding sites in the wwPDB based on metrics of regional structural quality with respect to the underlying electron density data used to derive the metal binding site structure. Our goal is to provide a set of metal binding sites that are of high regional quality for a wide variety of downstream structural, dynamic, and functional analyses.

## 2. Methods

Structural data from wwPDB listed on Jul 3, 2018 was used for the analysis. Their electron density data, if available, was acquired from the PDBe website. We used version 1.0 of the pdb-eda Python package [18] to analyze all downloaded PDB entries and matching electron density maps. Metal ions were detected across these PDB entries and filtered against four major quality control criteria:

1) electron density resolution less than or equal to 2.5 Å.
2) atom occupancy greater than or equal to 0.9.
3) no symmetry atoms within 3.5 Å.
4) sum of electrons of discrepancy within a 3.5 Å region surrounding the metal ion point position is less than a data-derived cutoff.

The resolution and occupancy information were retrieved directly from the PDB structure entry file in PDB Molecular Format (ent) format. We considered a resolution less than or equal to 2.5 Å and an occupancy greater than or equal to 0.9 as meeting an acceptable level of quality for most downstream structural and dynamic studies, since water and small ligands are typically visible below this resolution. Symmetry related atoms were calculated from the REMARK SMTRY records in the PDB structure data, as we took into account nearby asymmetric units. Atom-atom distance between a metal ion and all symmetry related atoms were computed and metal ions were filtered out if any symmetry atom point positions were present within 3.5 Å of the metal ion point position. Electron density maps were analyzed using the self-developed Python package, pdb-eda. This package provides methods for converting the electron density discrepancies in Fo-Fc density maps into numbers of electrons of discrepancy, when a significant protein component exists in the entry. A sum of the absolute value of both positive and negative electron discrepancies was computed for all significant discrepancies within 3.5 Å of the metal. Significant discrepancies were decided by a 3 standard deviation cutoff based on each individual electron density map, which is the commonly accepted cutoff for visualizing significant electron density map discrepancies. After filtering by criteria 1 and 2, we derived the distribution of all metal ion electron discrepancies sums and filtered out outliers by a 2 standard deviation cutoff. With the resulting filtered distribution, we set a max electron discrepancy sum cutoff to 1 standard deviation of this distribution. The electron density overlay graphs were prepared using the LiteMol Viewer [19]. All results and code used to generate the results for this study are available on FigShare: https://doi.org/10.6084/m9.figshare.8044451.

## 3. Results

We started with a list of 141,616 usable PDB entries, and 53,146 of them contains at least one metal ion. It is about 38 percent, which agrees very well with other studies and predictions [1,2]. In this study, we considered four major criteria in filtering “high-quality” metal ions, usable for downstream structural and dynamic analyses: resolution, occupancy, presence of symmetry atoms, and significant discrepancies in terms of numbers of electrons. Figure 1 shows both high-and low-quality examples based on these four criteria, as illustrated in an overlay of the electron density map over the structural model.

**Figure 1.**
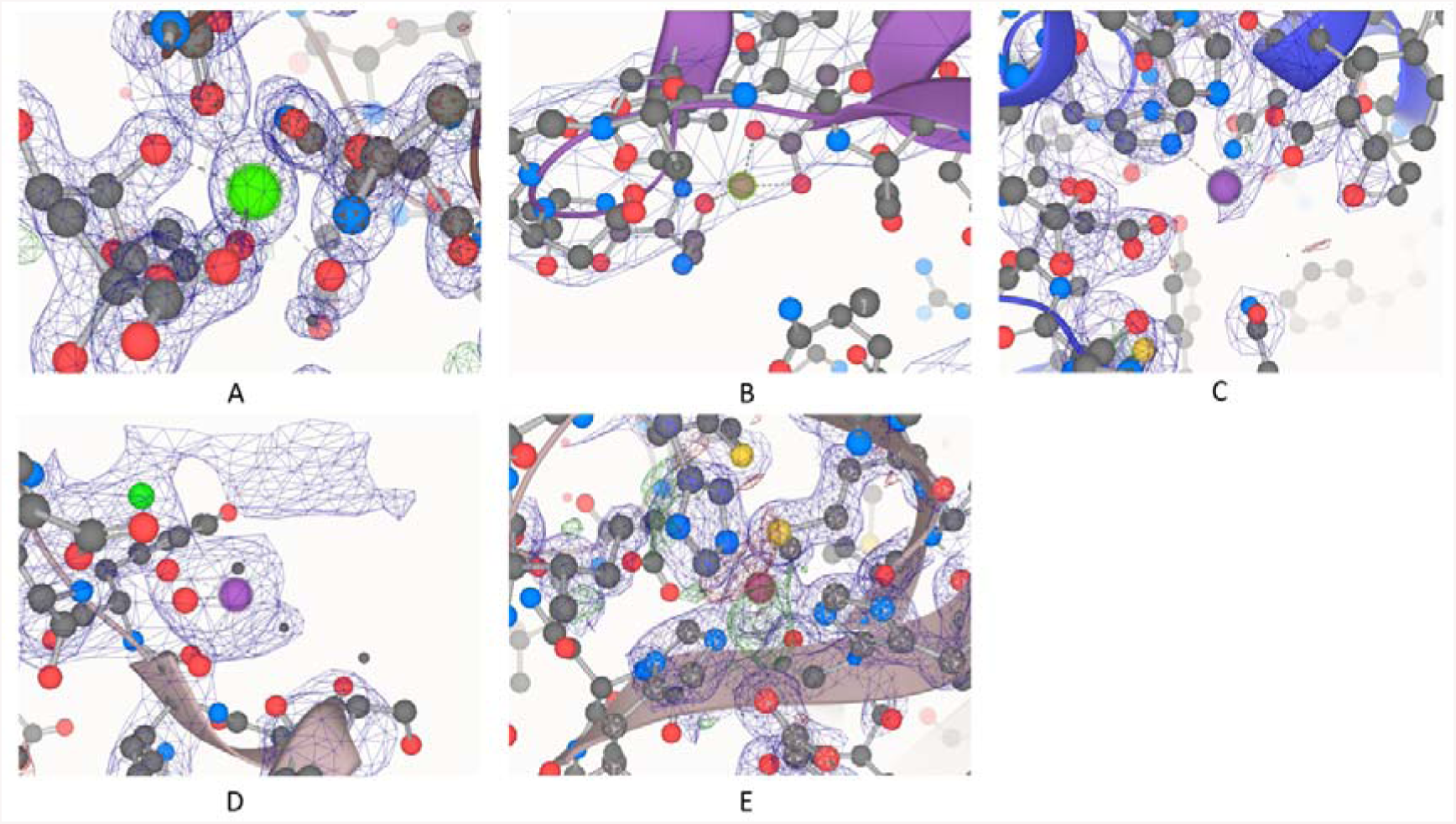
High and low quality examples for all four criteria: (A) PDB ID: 5FVN.F.405.CA, representative of high-quality by passing all four criteria; (B) PDB ID: 1YV0.C.163.MG, resolution: 7 Å; (C) PDB ID: 3CIA.B.701.ZN, occupancy: 0.7; (D) PDB ID: 5ER8.A.706.MN, symmetry atoms nearby; (E) PDB ID: 3LZQ.A.200.CU, high discrepancy between calculated and observed electron density maps.

Table 1 shows a tabulation of 56 different elemental types of metal ions observed in the wwPDB, with respect to four quality-control criteria. Zinc is present in the highest number of PDB entries, while magnesium has the highest number of metal sites. This is probably due to the presence of large numbers of magnesium ions in certain PDB entries like those of the ribosome [20]. Overall, 9 metals had over 1000 examples across the PDB that passed the four criteria. An additional 8 metals had over 100 examples that passed all four criteria. For the rest of this study, we will look at each of the four criteria in more detail, with respect to five of the most important and common metal ions in biochemistry: zinc, calcium, iron, manganese, and copper.

**Table 1.** Tabulation of different elemental types of metal ions observed in the wwPDB with respect to four quality-control criteria.

| Metal | Number of PDB entries | Number of total metal ion sites | Number with < 2.5 Å resolution | Number with high occupancy | Number with no nearby symmetry atoms | Number with no significant discrepancy densities | Number that passes all criterion |
| --- | --- | --- | --- | --- | --- | --- | --- |
| Zn | 13497 | 67405 | 56176 | 62550 | 62967 | 23883 | 13230 |
| Mg | 13225 | 85080 | 30537 | 81528 | 81463 | 48708 | 18305 |
| Ca | 10138 | 44538 | 36193 | 41929 | 41176 | 23836 | 15787 |
| Fe | 8054 | 41898 | 32283 | 39207 | 41499 | 20881 | 14427 |
| Na | 7516 | 23697 | 16645 | 22523 | 21072 | 18295 | 10700 |
| Mn | 3177 | 14347 | 9037 | 12089 | 13630 | 8755 | 4275 |
| K | 2390 | 8819 | 5671 | 7498 | 7678 | 5541 | 2973 |
| Ni | 1533 | 3578 | 2803 | 2752 | 2943 | 2093 | 986 |
| Cu | 1469 | 6918 | 5913 | 5548 | 6474 | 3676 | 2111 |
| Co | 1146 | 3601 | 2897 | 2920 | 3288 | 1687 | 976 |
| Cd | 926 | 6535 | 5351 | 4708 | 4863 | 2334 | 624 |
| Hg | 640 | 2302 | 1525 | 808 | 2223 | 528 | 11 |
| U | 507 | 6032 | 5522 | 4553 | 5196 | 2351 | 1693 |
| Pt | 242 | 869 | 564 | 212 | 802 | 249 | 4 |
| Mo | 209 | 785 | 685 | 505 | 692 | 323 | 147 |
| Al | 189 | 399 | 187 | 390 | 399 | 217 | 112 |
| Be | 187 | 510 | 273 | 461 | 504 | 318 | 175 |
| Ba | 166 | 900 | 399 | 558 | 733 | 186 | 6 |
| Ru | 162 | 341 | 288 | 163 | 318 | 113 | 8 |
| Sr | 151 | 3972 | 1394 | 3764 | 3869 | 2846 | 745 |
| V | 143 | 488 | 285 | 399 | 462 | 219 | 130 |
| Cs | 115 | 666 | 402 | 251 | 526 | 226 | 14 |
| W | 96 | 1743 | 396 | 1218 | 1639 | 280 | 15 |
| Yb | 91 | 247 | 189 | 136 | 127 | 60 | 6 |
| Au | 90 | 437 | 275 | 120 | 373 | 136 | 2 |
| Li | 73 | 124 | 110 | 116 | 109 | 96 | 72 |
| Gd | 65 | 444 | 408 | 268 | 409 | 134 | 22 |
| Pb | 62 | 229 | 113 | 87 | 187 | 63 | 5 |
| Y | 58 | 218 | 168 | 154 | 108 | 77 | 16 |
| Tl | 54 | 400 | 143 | 119 | 383 | 82 | 1 |
| Ir | 51 | 333 | 132 | 138 | 317 | 44 | 0 |
| Rb | 49 | 229 | 139 | 73 | 174 | 83 | 4 |
| Sm | 45 | 205 | 106 | 132 | 142 | 42 | 11 |
| Ag | 34 | 381 | 329 | 361 | 365 | 67 | 25 |
| Pr | 31 | 77 | 56 | 46 | 40 | 23 | 4 |
| Eu | 24 | 71 | 64 | 14 | 60 | 23 | 3 |
| Pd | 24 | 108 | 108 | 55 | 79 | 19 | 2 |
| Os | 23 | 101 | 34 | 77 | 97 | 20 | 3 |
| Re | 21 | 71 | 71 | 27 | 68 | 13 | 3 |
| Rh | 20 | 68 | 68 | 25 | 62 | 18 | 1 |
| Tb | 18 | 168 | 139 | 134 | 157 | 20 | 3 |
| Ta | 18 | 529 | 106 | 42 | 503 | 199 | 0 |
| Lu | 15 | 62 | 46 | 31 | 54 | 19 | 0 |
| Ho | 13 | 55 | 47 | 43 | 44 | 7 | 0 |
| La | 11 | 115 | 107 | 106 | 112 | 1 | 1 |
| Cr | 10 | 53 | 43 | 49 | 52 | 8 | 5 |
| Ga | 10 | 80 | 80 | 80 | 80 | 5 | 5 |
| Sn | 9 | 16 | 16 | 6 | 16 | 2 | 0 |
| Sb | 5 | 10 | 4 | 6 | 10 | 7 | 3 |
| Ce | 4 | 70 | 70 | 66 | 70 | 0 | 0 |
| Er | 3 | 18 | 0 | 17 | 0 | 1 | 0 |
| Zr | 3 | 31 | 28 | 30 | 0 | 0 | 0 |
| In | 2 | 3 | 1 | 3 | 0 | 1 | 0 |
| Bi | 2 | 2 | 2 | 0 | 2 | 0 | 0 |
| Hf | 2 | 44 | 44 | 43 | 0 | 10 | 9 |
| Dy | 1 | 26 | 26 | 0 | 0 | 18 | 0 |
| Total | 66819 | 330448 | 218698 | 299138 | 308616 | 168843 | 87660 |

### 3.1. Resolution

The electron density resolution represents an overall metric of structural quality for an x-ray crystallographic structure. Structural entries with a resolution below 2.5 Å are generally considered usable for many structural and dynamics analyses. Figure 2 illustrates the distribution of resolution for the top five most essential metals in biology. In general, the overall and individual metal ions have similar distributions. The distribution for manganese has several spikes, which is mainly due to the over-representation of replicate values from structural entries with large numbers of manganese ions.

**Figure 2.**
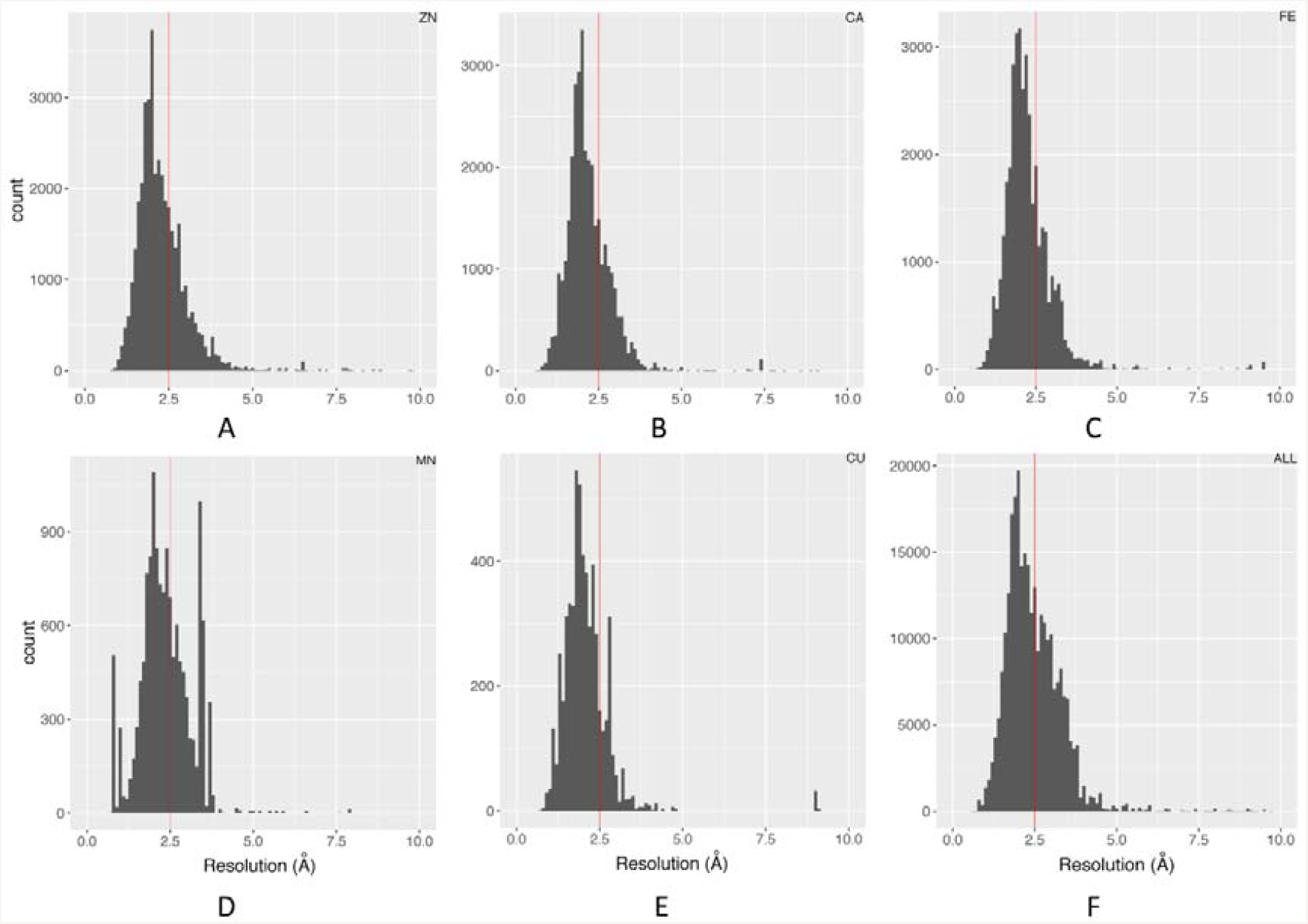
Distribution and 2.5 Å cutoff of structure resolutions: (A) Zn; (B) Ca; (C) Fe; (D) Mn; (E) Cu; (F) All metal ions.

### 3.2. Occupancy and Symmetry related atoms

The majority of metal ions have an occupancy of 1. However, there are two general cases where low occupancy occurs. In the first general case, when there is more than one conformation available during the structure determination, multiple conformations (typically two) are often kept in the data and are often marked as “ALT”. Thus, different conformations will only possess the metal ions with partial occupancy. Typically for two conformations, the occupancy will be 0.5 for each conformation. For the second general case, only a fraction of the repeating unit cells in the protein crystal has the observed metal ion, and the occupancy will represent the percentage of the crystal structure with the metal ion bound. In either case, low occupancy sites can be considered low-quality for aggregated analyses, since only a fraction of the experimental data supports the given model of the metal ion position. A filter of 0.9 occupancy removes about 10% of all metal ions and is consistent for most of individual metal ions. Therefore, this criterion only removes a minority of metal ion sites.

Crystal contacts can pose as an artifact, affecting the binding of the metal ion, especially on the surface of a protein structure. It may represent a false binding that does not biologically exist, i.e. when the crystal packing environment is no longer available. Also, crystal contacts can affect protein-ligand binding [14]. Our study demonstrates that only about 7% of metal ion sites are filtered out by a 3.5 Å symmetry atom exclusion criterion.

### 3.3. Discrepancy between calculated and observed electron density maps

Figure 1 demonstrates why electron density maps can be extremely useful for validating high-quality regions within protein structures. As described in the methods, we used our pdb-eda Python package to compare every metal binding site to its Fo-Fc electron density map. Figure 3 shows the distributions of the sum of absolute electron disagreement within a 3.5 Å radius of each metal ion. Overall and individual metal ions demonstrate very similar distributions, justifying the calculation of a single data driven cutoff from the overall distribution. The final data-driven cutoff for the sum of absolute electron discrepancy is approximately 19.3 electrons and is based on one standard deviation of the Figure 3F distribution with outliers removed. This is a purposely conservation quality control criterion, representing roughly two water molecules worth of electron discrepancy. However, only 24.8% of metal ion sites are filtered out by this criterion.

**Figure 3.**
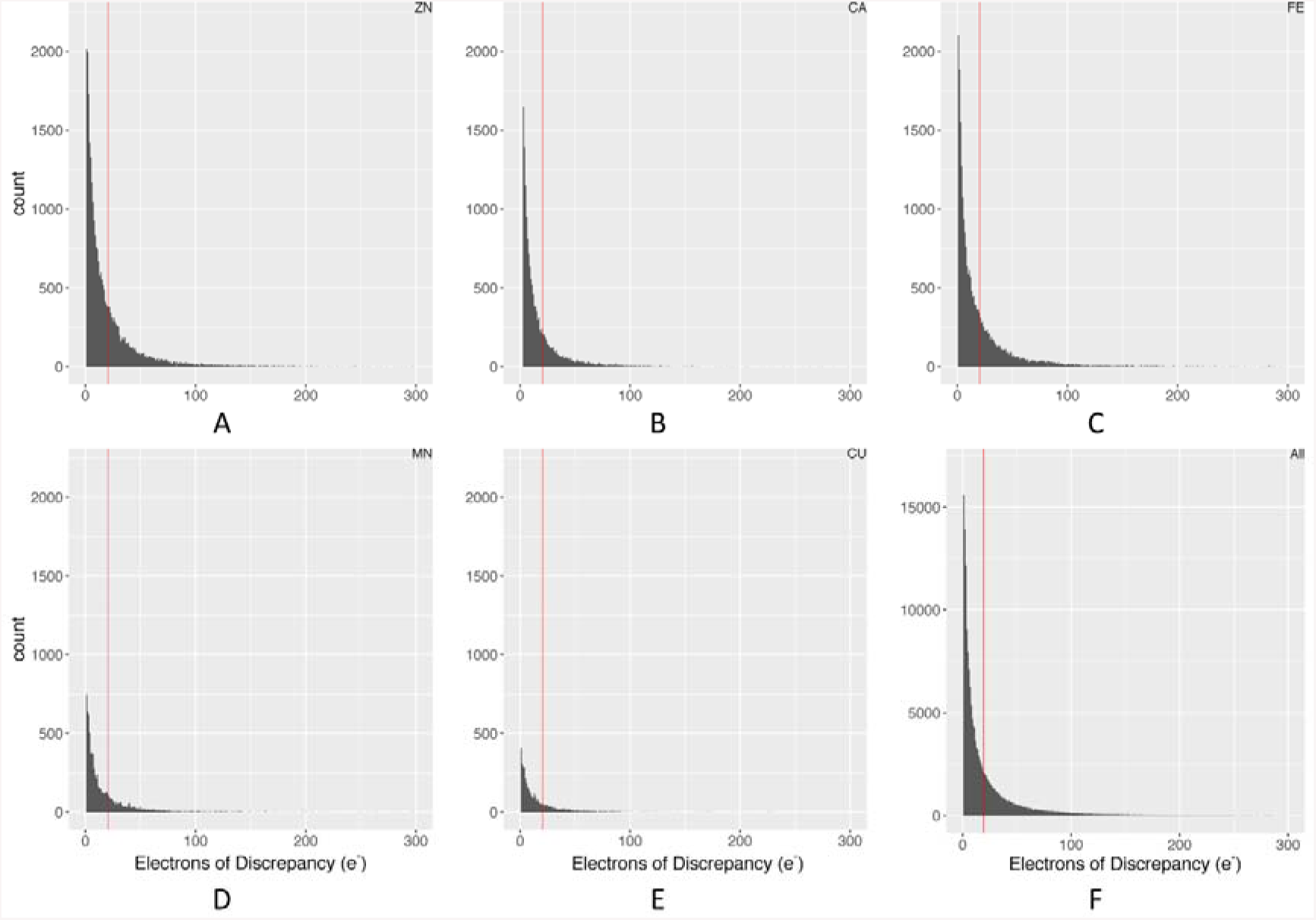
Distribution of electron discrepancy within 3.5 Å of the metal ion: (A) Zn; (B) Ca; (C) Fe; (D) Mn; (E) Cu; (F) All metal ions. The red line represents a 1 standard deviation cutoff calculated from the distribution in graph F with outliers removed.

## 4. Discussion

As illustrated by previous studies, regional structural quality affects the usability of bound ligand structure, including bound metal ions, for accurately interpreting structural, dynamic, and chemical properties of ligand binding sites [14,21,22]. Therefore, steps should be taken to ensure the quality of metal binding sites for various downstream structural and dynamics analyses. As the number of structures available in PDB continues to grow dramatically every year, more attention is being paid to ensuring that only high-quality datasets are used in these studies. Toward this goal, we have developed new methods in the open-source pdb-eda Python package that enable the evaluation of regional structural quality with respect to underlying experimental data.

In this study, we demonstrate the use of electron density maps for a systematic evaluation of regional structural quality around all metal binding sites in the PDB with matching electron density maps provided by the PDBe. This is one of 4 criteria used for evaluating the structural quality of metal binding sites for the purpose of generating large high-quality datasets for downstream analyses. The other criteria include electron density resolution, exclusion of nearby symmetry atoms, and the occupancy of the metal ion. For commonly-bound metal ions, additional criteria can be used for quality control [13], including the evaluation of bond lengths between ligating atoms (using the coordination chemistry definition of ligand) and the metal ion and coordination shell considerations [8,23,24]. However, several of these evaluations require the accurate identification of ligating atoms and is complicated by the wide variation in coordination geometries. Moreover, these additional criteria cannot be practically applied to all 56 elemental types of metals analyzed in this study, given the current examples available in the PDB. Therefore, we limited to four criteria that could be straight-forwardly applied to all elemental types of metal ions currently present in the PDB.

A full list of metal binding sites that pass all four criteria utilized in this study can be found in Table S1 (in FigShare repository), along with the values used for criterion evaluation. In addition, all code used to generate this table is available in a FigShare repository along with all metals binding sites evaluated in this study. Therefore, metal binding sites can be re-evaluated against modified criteria that match the quality-control requirements of a given downstream analysis. Also with this code, future versions of the PDB can be analyzed in a similar manner to regenerate an updated list of metal binding sites.

In conclusion, we have demonstrated an approach for the pan-PDB evaluation of metal binding site structural quality with respect to underlying x-ray crystallographic experimental data represented in available electron density maps of proteins. Especially for non-crystallographers, we hope to change the focus and discussion of structural quality from a global evaluation to a regional evaluation, since all structural entries in the wwPDB appear to have both regions of high and low structural quality.

## Supplementary Materials

All results and code used to generate the results for this study are available on FigShare: https://doi.org/10.6084/m9.figshare.8044451.

## Author Contributions

S.Y. and H.M. conceptualized the study; S.Y. conducted the data analysis and visualization; S.Y. and H.M. validated the results, wrote the original draft, and edited and reviewed the manuscript.

## Acknowledgments

In the interest of full disclosure, the authors of this paper are not card-carrying crystallographers. However, we are conscientious biophysical informaticists.

## References

1. Andreini, C.; Bertini, I.; Rosato, A. Metalloproteomes: a bioinformatic approach. Acc Chem Res 2009, 42, 1471–1479, doi:10.1021/ar900015x.

2. Putignano, V.; Rosato, A.; Banci, L.; Andreini, C. MetalPDB in 2018: a database of metal sites in biological macromolecular structures. Nucleic Acids Res 2018, 46, D459–D464, doi:10.1093/nar/gkx989.

3. Ruta, L.L.; Nicolau, I.; Popa, C.V.; Farcasanu, I.C. Manganese Suppresses the Haploinsufficiency of Heterozygous trpy1Delta/TRPY1 Saccharomyces cerevisiae Cells and Stimulates the TRPY1-Dependent Release of Vacuolar Ca(2+) under H(2)O(2) Stress. Cells 2019, 8, doi:10.3390/cells8020079.

4. Sibarov, D.A.; Antonov, S.M. Calcium-Dependent Desensitization of NMDA Receptors. Biochemistry (Mosc) 2018, 83, 1173–1183, doi:10.1134/S0006297918100036.

5. Sievers, Q.L.; Petzold, G.; Bunker, R.D.; Renneville, A.; Slabicki, M.; Liddicoat, B.J.; Abdulrahman, W.; Mikkelsen, T.; Ebert, B.L.; Thoma, N.H. Defining the human C2H2 zinc finger degrome targeted by thalidomide analogs through CRBN. Science 2018, 362, doi:10.1126/science.aat0572.

6. Bhim, A.; Laha, S.; Gopalakrishnan, J.; Natarajan, S. Color Tuning in Garnet Oxides: The Role of Tetrahedral Coordination Geometry for 3 d Metal Ions and Ligand-Metal Charge Transfer (Band-Gap Manipulation). Chem Asian J 2017, 12, 2734–2743, doi:10.1002/asia.201701040.

7. Donahue, C.M.; McCollom, S.P.; Forrest, C.M.; Blake, A.V.; Bellott, B.J.; Keith, J.M.; Daly, S.R. Correction to Impact of Coordination Geometry, Bite Angle, and Trans Influence on Metal-Ligand Covalency in Phenyl-Substituted Phosphine Complexes of Ni and Pd. Inorg Chem 2015, 54, 8857, doi:10.1021/acs.inorgchem.5b01794.

8. Yao, S.; Flight, R.M.; Rouchka, E.C.; Moseley, H.N. Aberrant coordination geometries discovered in the most abundant metalloproteins. Proteins 2017, 85, 885–907, doi:10.1002/prot.25257.

9. Gil-Moreno, S.; Jimenez-Marti, E.; Palacios, O.; Zerbe, O.; Dallinger, R.; Capdevila, M.; Atrian, S. Does Variation of the Inter-Domain Linker Sequence Modulate the Metal Binding Behaviour of Helix pomatia Cd-Metallothionein? Int J Mol Sci 2015, 17, doi:10.3390/ijms17010006.

10. M’Kandawire, E.; Mierek-Adamska, A.; Sturzenbaum, S.R.; Choongo, K.; Yabe, J.; Mwase, M.; Saasa, N.; Blindauer, C.A. Metallothionein from Wild Populations of the African Catfish Clarias gariepinus: From Sequence, Protein Expression and Metal Binding Properties to Transcriptional Biomarker of Metal Pollution. Int J Mol Sci 2017, 18, doi:10.3390/ijms18071548.

11. Warren, G.L.; Do, T.D.; Kelley, B.P.; Nicholls, A.; Warren, S.D. Essential considerations for using protein-ligand structures in drug discovery. Drug Discov Today 2012, 17, 1270–1281, doi:10.1016/j.drudis.2012.06.011.

12. Berman, H.; Henrick, K.; Nakamura, H.; Markley, J.L. The worldwide Protein Data Bank (wwPDB): ensuring a single, uniform archive of PDB data. Nucleic Acids Res 2007, 35, D301–303, doi:10.1093/nar/gkl971.

13. Zheng, H.; Cooper, D.R.; Porebski, P.J.; Shabalin, I.G.; Handing, K.B.; Minor, W. CheckMyMetal: a macromolecular metal-binding validation tool. Acta Crystallogr D Struct Biol 2017, 73, 223–233, doi:10.1107/S2059798317001061.

14. Sondergaard, C.R.; Garrett, A.E.; Carstensen, T.; Pollastri, G.; Nielsen, J.E. Structural artifacts in protein-ligand X-ray structures: implications for the development of docking scoring functions. J Med Chem 2009, 52, 5673–5684, doi:10.1021/jm8016464.

15. Smart, O.S.; Horsky, V.; Gore, S.; Svobodova Varekova, R.; Bendova, V.; Kleywegt, G.J.; Velankar, S. Validation of ligands in macromolecular structures determined by X-ray crystallography. Acta Crystallogr D Struct Biol 2018, 74, 228–236, doi:10.1107/S2059798318002541.

16. Kleywegt, G.J.; Harris, M.R.; Zou, J.Y.; Taylor, T.C.; Wahlby, A.; Jones, T.A. The Uppsala Electron-Density Server. Acta Crystallogr D Biol Crystallogr 2004, 60, 2240–2249, doi:10.1107/S0907444904013253.

17. Gutmanas, A.; Alhroub, Y.; Battle, G.M.; Berrisford, J.M.; Bochet, E.; Conroy, M.J.; Dana, J.M.; Fernandez Montecelo, M.A.; van Ginkel, G.; Gore, S.P., et al. PDBe: Protein Data Bank in Europe. Nucleic Acids Res 2014, 42, D285–291, doi:10.1093/nar/gkt1180.

18. Yao, S.; Moseley, H.N.B. A chemical interpretation of protein electron density maps in the worldwide protein data bank. bioRxiv 2019, 10.1101/613109, 613109, doi:10.1101/613109.

19. Sehnal, D.; Deshpande, M.; Varekova, R.S.; Mir, S.; Berka, K.; Midlik, A.; Pravda, L.; Velankar, S.; Koca, J. LiteMol suite: interactive web-based visualization of large-scale macromolecular structure data. Nat Methods 2017, 14, 1121–1122, doi:10.1038/nmeth.4499.

20. Nierhaus, K.H. Mg2+, K+, and the Ribosome. Journal of Bacteriology 2014, 196, 3817–3819, doi:10.1128/jb.02297-14.

21. Tang, Y.T.; Marshall, G.R. PHOENIX: a scoring function for affinity prediction derived using high-resolution crystal structures and calorimetry measurements. J Chem Inf Model 2011, 51, 214–228, doi:10.1021/ci100257s.

22. Wang, R.; Lai, L.; Wang, S. Further development and validation of empirical scoring functions for structure-based binding affinity prediction. J Comput Aided Mol Des 2002, 16, 11–26.

23. Yao, S.; Flight, R.M.; Rouchka, E.C.; Moseley, H.N. A less biased analysis of metalloproteins reveals novel zinc coordination geometries. Proteins: Structure, Function, and Bioinformatics 2015, 10.1002/prot.24834, doi:10.1002/prot.24834.

24. Yao, S.; Flight, R.M.; Rouchka, E.C.; Moseley, H.N. Perspectives and expectations in structural bioinformatics of metalloproteins. Proteins: Structure, Function, and Bioinformatics 2017, 85, 938–944.

